# Protocol for 3D digital dynamic histomorphometry of bone via time-lapse registration of serial microCT scans

**DOI:** 10.64898/2026.05.28.728585

**Authors:** Quentin A. Meslier, Nicole Migotsky, Syeda N. Lamia, Erica L. Scheller, Matthew J. Silva

## Abstract

Serial *in vivo* microCT enables quantification of dynamic bone remodeling by capturing formation and resorption over time. This protocol uses Dragonfly to register pre- and post-intervention scans of rodent long bones through an intuitive drag-and-click workflow that requires no coding expertise. Rigid registration enables spatial mapping and quantification of formed, resorbed, and quiescent cortical and trabecular bone at periosteal and endosteal surfaces across multiple tibial regions, providing a non-destructive and complementary alternative to classic, fluorochrome-based dynamic histomorphometry.

**Graphical abstract:** 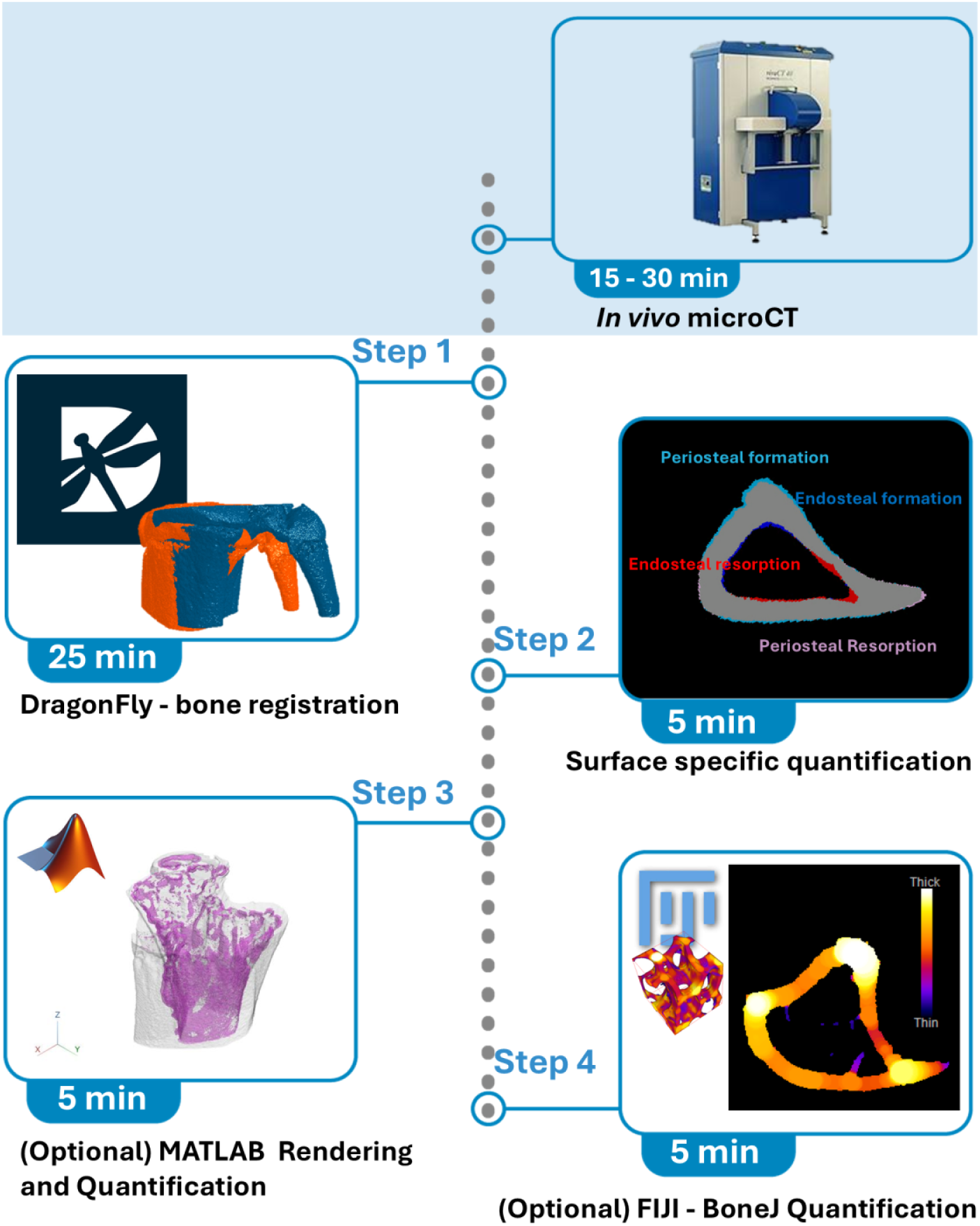

## Introduction

### Innovation

Traditional approaches for assessing bone remodeling rely on *ex vivo* microCT or histomorphometry, with complementary histological staining of bone cells^1,2^. *Ex vivo* microCT captures bone structure at a single endpoint, while classic dynamic histomorphometry with serial fluorochrome labels is used to quantify bone formation but does not allow for quantitative analysis of bone resorption. Histological staining to identify osteoclasts (e.g., by tartrate-resistant acid phosphatase (TRAP)) or eroded surface can provide insights into resorptive activity but does not provide a measure the amount of bone resorbed in a time interval. As a result, resolving both sides of dynamic bone (re)modeling, i.e., bone formation and bone resorption, in the same specimen over time has not been feasible

Enabled by *in vivo* microCT, methods to overcome these limitations have been developed ^3–5^. These methods provide a complementary approach to traditional methods by enabling registration (overlay) of serial *in vivo* microCT scans to simultaneously quantify bone formation, resorption, and quiescence throughout the course of an experiment. The protocol presented herein provides a step-by-step pipeline for users to perform these analyses. By aligning pre-intervention and post-intervention scan datasets, the method captures time-resolved, spatially mapped (re)modeling events at periosteal and endosteal surfaces, offering a form of “3D digital dynamic histomorphometry” derived from non-destructive imaging.

A key innovation herein is the accessibility of the registration workflow. Previously reported time-lapse microCT registration techniques require custom scripts and advanced coding expertise, limiting adoption ^3,5^. In contrast, this protocol uses Dragonfly interactive software to provide a beginner-friendly, drag-and-click interface that eliminates the need for programming while preserving analytical rigor. This lowers the technical barrier for researchers and broadens access to dynamic bone remodeling metrics that were previously restricted to specialized computational pipelines.

By merging *in vivo* microCT with intuitive image registration, this protocol delivers a practical and scalable approach for capturing dynamic bone biology, complementing and extending traditional histomorphometric analyses.

Using this approach, we were recently able to evaluate changes in load-induced bone remodeling in mice lacking the citrate transporter *Slc13a5* in osteoblasts compared with controls which further supports the applicability of this protocol beyond the validation data presented here.

### Institutional permissions

All animal experiments were performed in accordance with institutional guidelines and with prior approval from the Washington University Institutional Animal Care and Use Committee (IACUC). Female C57BL/6J mice (Jackson Laboratories) were group-housed in the Washington University animal facility under controlled conditions (22–25 °C, 12 h light/dark cycle) with ad libitum access to water and a standard rodent diet (Purina 5053; Purina, St. Louis, MO, USA).

Researchers intending to perform similar experiments should obtain appropriate institutional animal care and use approval before initiating any procedures.

### Preparation: *In vivo* scanning

#### Timing

∼15 min per scan (213 slices, voxel size: 10.5 µm) on a Scanco vivaCT 40; Total time varies depending on scanner model, resolution, and size of the region-of-interest (ROI). The scanning parameters used in this protocol were: 10.5 μm/voxel, 70 kVp, 114 μA, 300 ms integration.

1. Acquire *in vivo* microCT scans of the target bone (e.g., tibia) prior to the experimental intervention (Pre-Scan) and repeat scanning at the experimental endpoint (Post-Scan) using identical scan parameters
2. Pre- and Post-scans should cover the same bone region. Use a landmark to guarantee that the same region is being scanned (growth plate, tibia-fibula junctions, etc).
3. Visually inspect reconstructed images immediately after acquisition to confirm the absence of motion artifacts, blurring, or slice misalignment
4. Export all validated scans in DICOM format for subsequent image registration and analysis.

##### CRITICAL

Motion artifacts and image smearing significantly reduce registration accuracy and will compromise quantification of formed and resorbed bone. The scan should be repeated if images present significant artifacts. In addition, Pre- and Post-scans need to overlap enough to allow registration of the area of interest to result in meaningful quantification.

## Key resources table

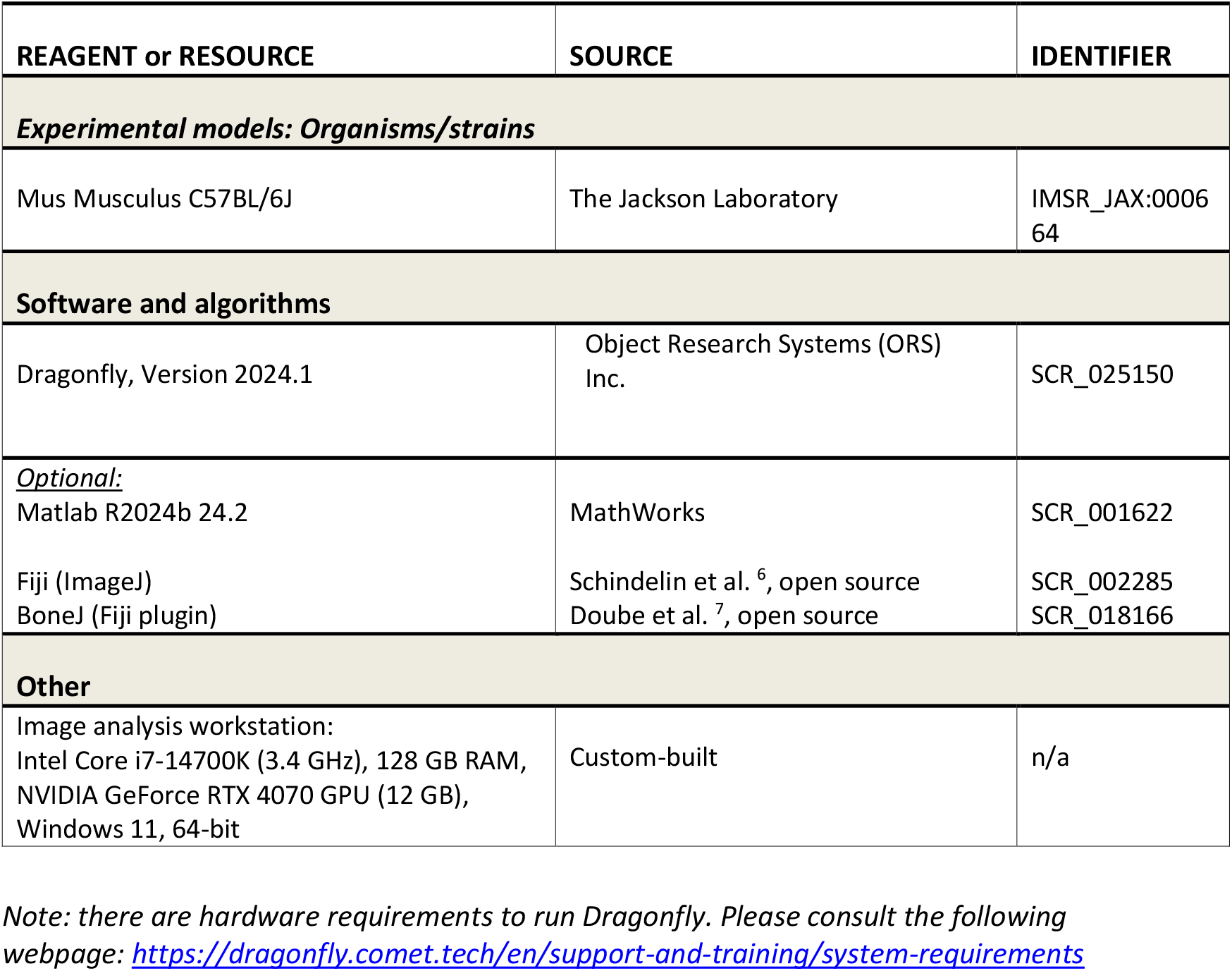

| REAGENT or RESOURCE | SOURCE | IDENTIFIER |
| --- | --- | --- |
| ***Experimental models: Organisms/strains*** |  |  |
| Mus Musculus C57BL/6J | The Jackson Laboratory | IMSR_JAX:000664 |
| **Software and algorithms** |  |  |
| Dragonfly, Version 2024.1 | Object Research Systems (ORS) Inc. | SCR_025150 |
| *Optional:*<br>Matlab R2024b 24.2 | MathWorks | SCR_001622 |
| Fiji (ImageJ)<br>BoneJ (Fiji plugin) | Schindelin et al. <sup>6</sup>, open source<br>Doubé et al. <sup>7</sup>, open source | SCR_002285<br>SCR_018166 |
| **Other** |  |  |
| Image analysis workstation:<br>Intel Core i7-14700K (3.4 GHz), 128 GB RAM,<br>NVIDIA GeForce RTX 4070 GPU (12 GB),<br>Windows 11, 64-bit | Custom-built | n/a |

*Note: there are hardware requirements to run Dragonfly. Please consult the following webpage: https://dragonfly.comet.tech/en/support-and-training/system-requirements*

## Methods

In this step-by-step section, we used Pre- and Post-scans from a mouse tibia midshaft.

### Load scans in Dragonfly

This first step will walk you through the upload of the Pre- and Post-scans into Dragonfly.

#### Timing: 5 min

1. Select “File” and “Import DICOM Images”
2. Click on “Folder Content Tab”
3. Click on “Open Folder”
4. Browse for folder of DICOM images
5. Click “Ok”
6. Click on the Scan info to highlight it
7. Click on “View Study”
8. Rename data set as “PreScan_ScanNbr”
  a. Double click on the data set
  b. Rename and adapt the “ScanNbr” based on your experiment
9. Repeat for PostScan data set
  a. Repeat steps 1 to 8
10. Scans should now appear in the Data properties and Settings section (Fig.1)

**Figure 1:**
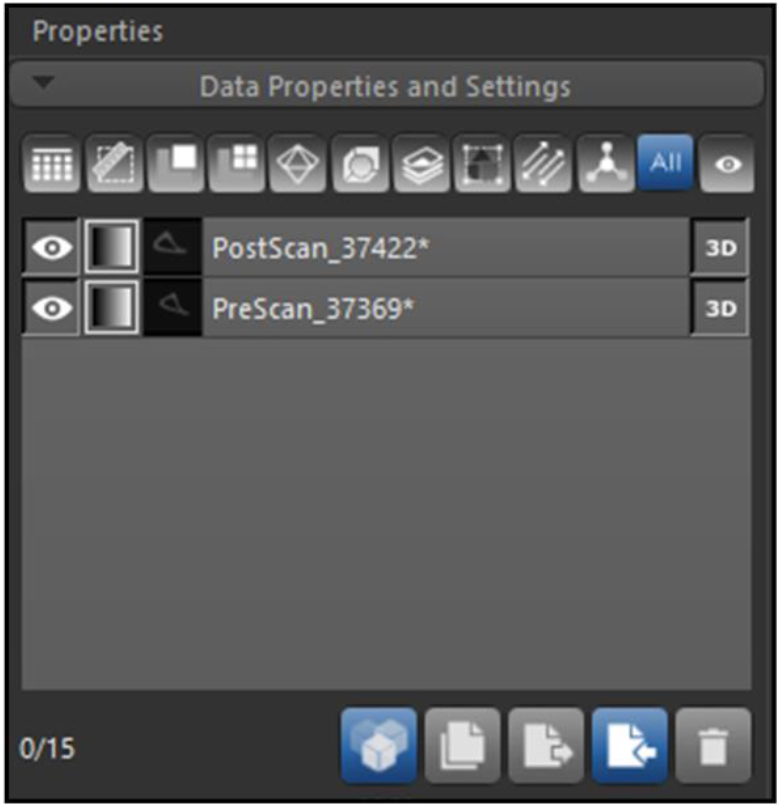
“Data properties and Settings” window showing the uploaded Pre- and Post-scans in Dragonfly.

*Note: Save ORS session regularly*

### Create masks and remove scans background

#### Timing: 5 min

This step removes the background of the scans to focus only on the bone and facilitates the registration.

11. Make PreScan_ScanNbr visible
  a. Click on the eye icon next to the data set name
  b. Click on the data set name to highlight it

*Note: Make sure no other data set is visible*

12. Create PreScan_ScanNbr mask
  a. Go to the “Segment Tab”
  b. Check “Define Range” in the “Range” menu
  c. Click “Upper Otsu”. Only the bone should show up red *Note: See Troubleshooting 1* *Note: Make sure not to click multiple times on Upper Ostu. If you do, click “Reset under “Selected Range” to reset the defined range*
  d. Record the Upper Otsu numbers for future reference
  e. Click “Add to New” to add the mask to the “Data Properties and settings” tab
  f. Unclick “Define range”
  g. Rename the data set that was just created (default is New ROI)
    1. Rename it to “PreROI”
  h. Select PreROI to highlight it
  i. Set dilation parameters under “Morphological operations” (Fig.2)
    1. Dimensionality: Select “2D (Z)”
    2. Shape: Select “Circle”
    3. Kernel Size: Select “15” *Note: for proximal tibia registration use a kernel size of 11*
  j. Click “Dilate”
  k. Click “Smooth”
  l. Right Click on PreROI, click “Modify and Transform” and Select “Invert”
    1. Check “Invert Values”
    2. Check “Create New DataSet”
    3. Click “Apply”
    4. Click “Close” *Note: the inverted ROI will be named PreROI(Inverted). The background will be masked. The bone should not be. This is where manual painting can be done to ensure all bone pixels are NOT masked and all the background and noise IS masked*.
  m. Keep PreROI(Inverted) selected
  n. Under “Image Operations”
    1. Select PreScan_ScanNbr in the drop-down menu
    2. Click “Overwrite”
    3. Enter “-3000” *Note: Depending on the pixel intensity of your background, you might need to enter a different overwriting value. We recommend entering the lowest number allowed (up to -20,000 if needed)*.
    4. Click “OK” *Note: if image artifacts are visible, they can be removed using the ROI painter (See Troubleshooting 1)*
13. Repeat step 11 for PostScan_ScanNbr
  a. Adjust names to PostROI and PostROI(Inverted) instead of PreROI and PreROI(Inverted)

**Figure 2:**
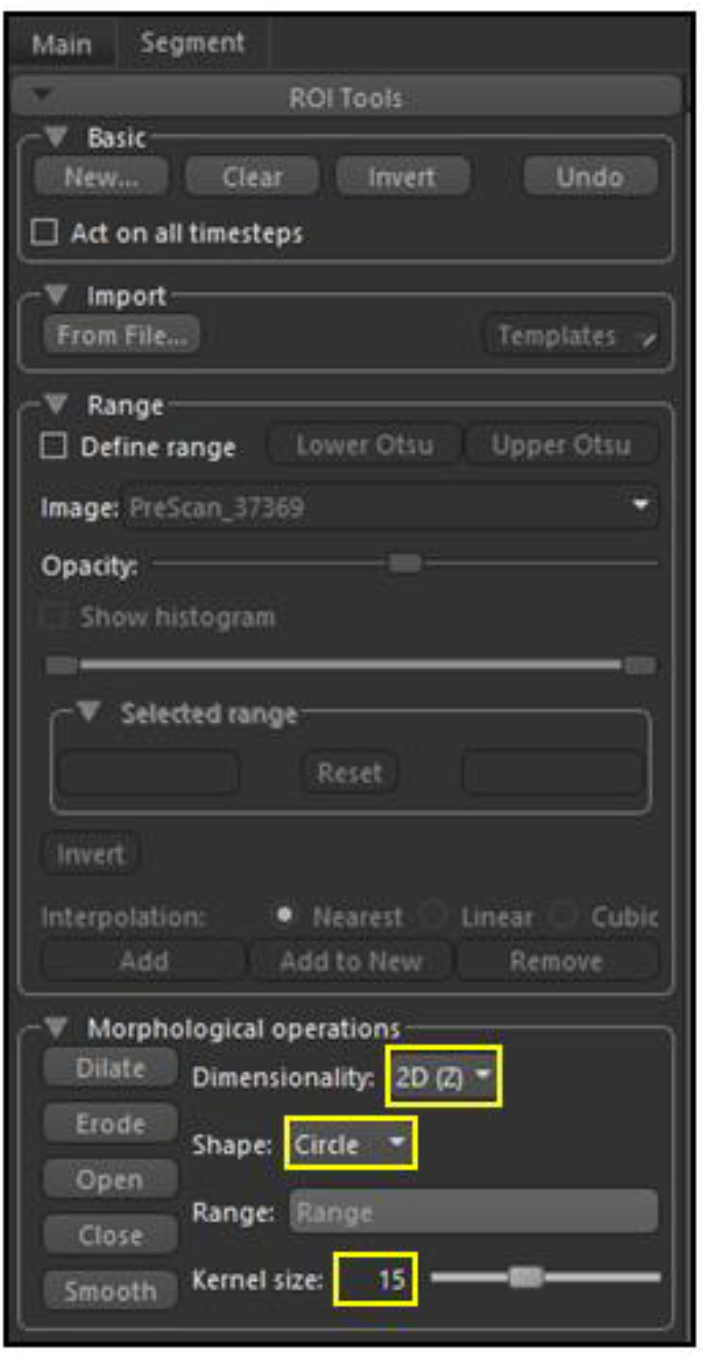
“Segment” menu showing the parameters set up for dilation and smoothing of the masks

### Align Centroids

#### Timing: 1 min

This step allows for an approximate alignment of two scans to facilitate image registration

14. Align centroid. This step aligns the centroid of the Pre-Scan and Post-Scan to facilitate the registration
  a. Right Click on PostScan_ScanNbr data set
  b. Select “Align” and “Centroid With …”
  c. Seletect PreScan_ScanNbr
  d. Click “OK”

*Note: See Troubleshooting 2*

*Note: After this step PreScan and PostScan now have their centroid aligned but they are not registered yet (Fig.4)*.

**Figure 3:**
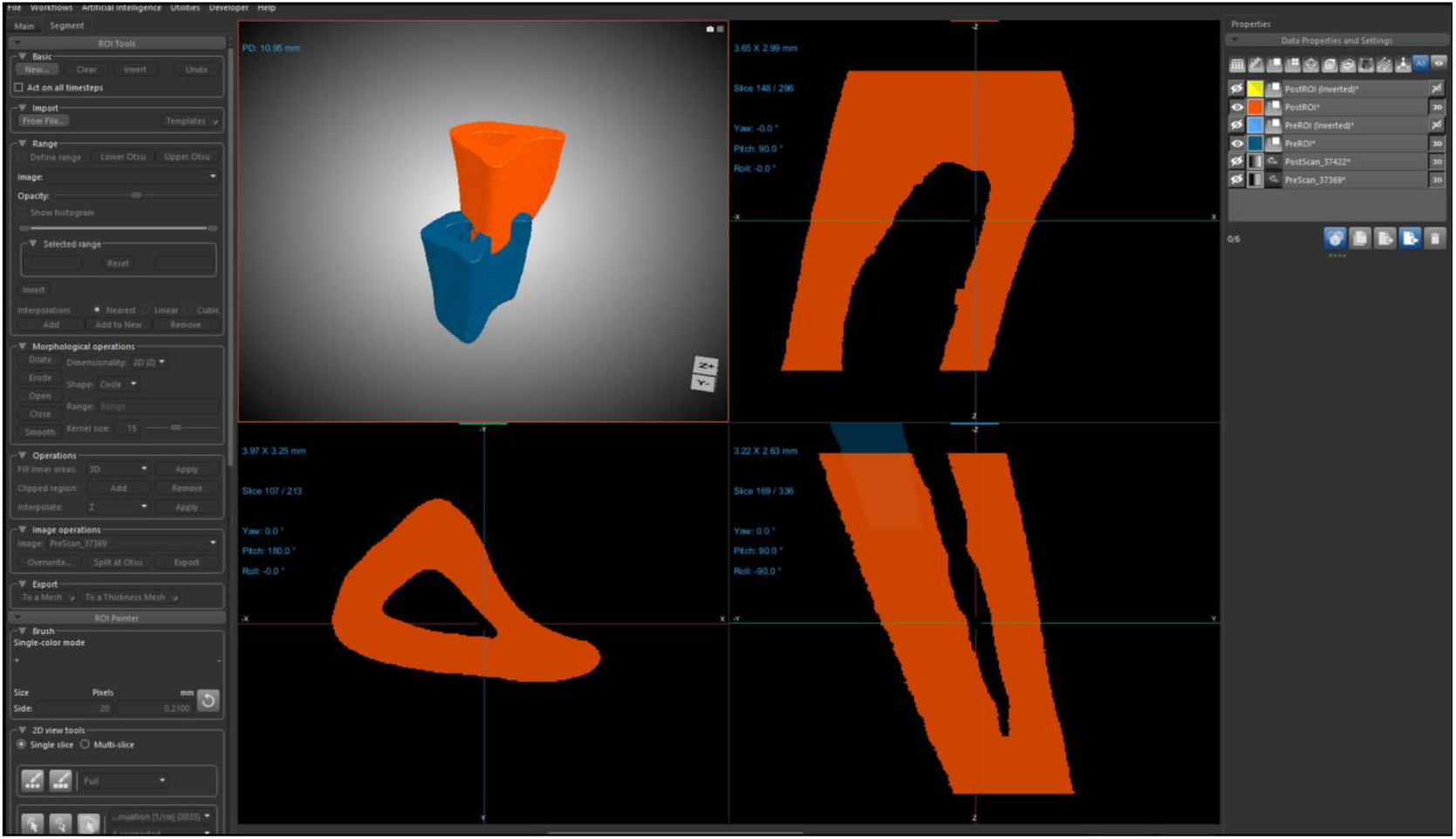
Dragonfly view of the two unregistered masks PreROI and PostROI

**Figure 4:**
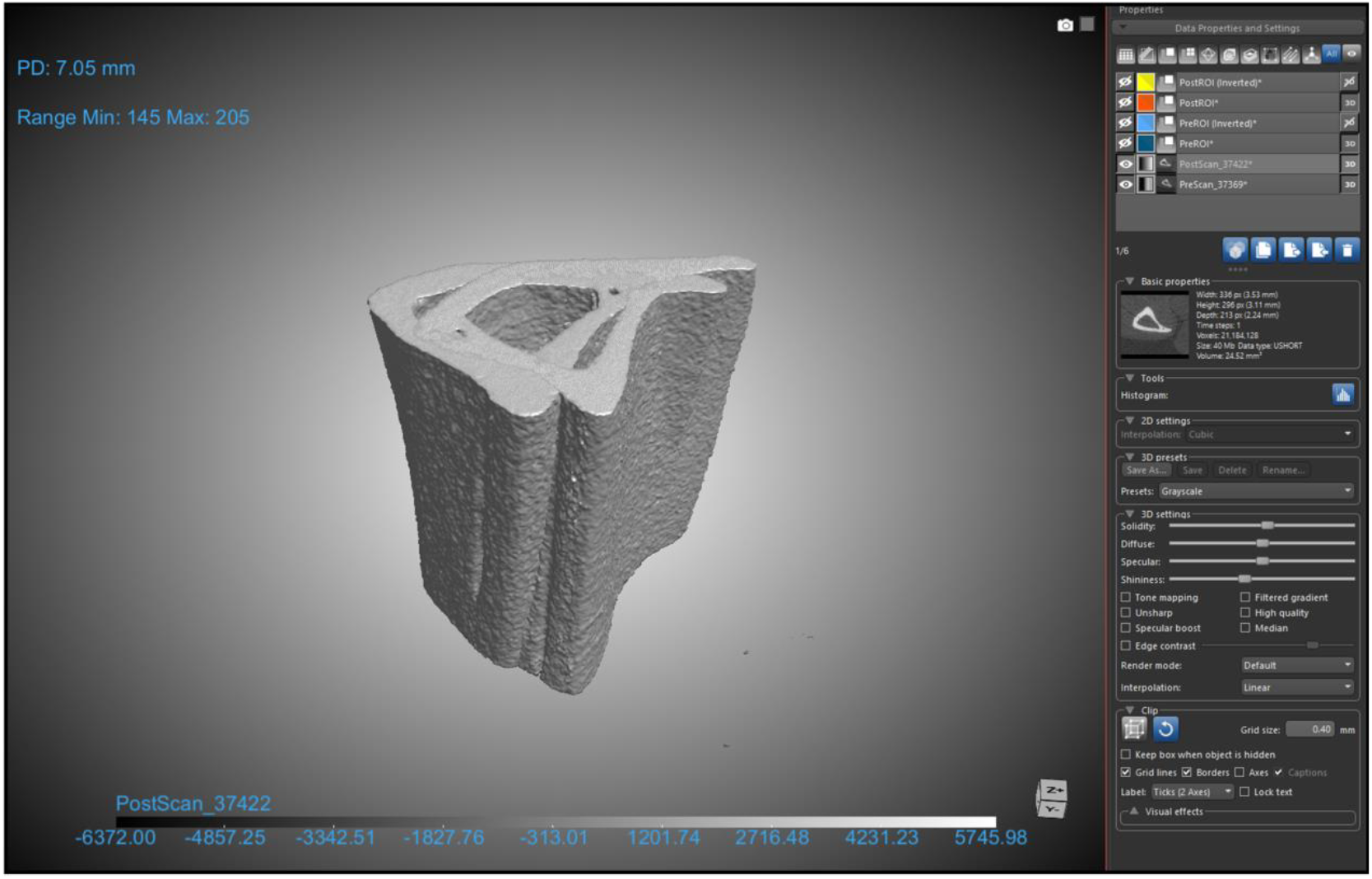
Dragonfly view of PreScan and PostScan centroids alignment

### Pre and Post scans registration

#### Timing: 10 min

This step allows for accurate registration of the Pre- and Post-scans.

15. Image registration
  a. Go to “Utilities” -> “Open Plugins” -> “Image Registration”. A window will open
  b. Adjust the parameter of the image registration
    1. Fixed: “PreScan_ScanNbr”
    2. Mobile: “PostScan_ScanNbr”
    3. Register using: “Rotation, Translation”
    4. Interpolation: “Linear”
    5. Method: “Mutual information”
  c. Start Coarse registration
    1. Enter the rotation and translation parameter as shown in Table 1. *Note: See Troubleshooting 3*
  d. Click “Apply”
  e. Once the registration step is completed, check the “Registration information section” (Fig.5A). Repeat Coarse registration until “Mutual Info” are similar on both side of the arrow, and until “Rotation and translation” are close or equal to 0. (Fig.5B) *Note: “Rotation” and “translation” values may not reach exactly zero in all data sets, but they should remain small and stable across repeated registrations, reflecting consistent and accurate alignment*.
  f. Run intermediate Registration
    1. Enter the parameters shown in Table 2 and click “apply”
  g. Repeat Intermediate registration until “Mutual Info “are similar on both side of the arrow, “Rotation” and “translation” are close or equal to 0.
  h. Start Fine Registration
    1. Enter the parameters shown in Table 3 and click “apply”
  i. Repeat Fine registration until “Mutual Info “are similar on both side of the arrow, “Rotation” and “translation” are close or equal to 0.
  j. PreScan and PostScan are now registered as shown in Fig.6

**Table 1:** Coarse Registration parameters.

| Registration | Initial step |  | Smallest step |
| --- | --- | --- | --- |
| Coarse registration | Rotation | 10 | 0.5 |
|  | Translation | 0.15 | 0.011 |

**Table 2:** Intermediate Registration parameters.

| Registration | Initial step |  | Smallest step |
| --- | --- | --- | --- |
| Intermediate registration | Rotation | 5 | 0.01 |
|  | Translation | 0.1 | 0.01 |

**Table 3:** Fine Registration parameters.

| Registration | Initial step |  | Smallest step |
| --- | --- | --- | --- |
| Fine Registration | Rotation | 5 | 0.01 |
|  | Translation | 0.1 | 0.001 |

**Figure 5:**
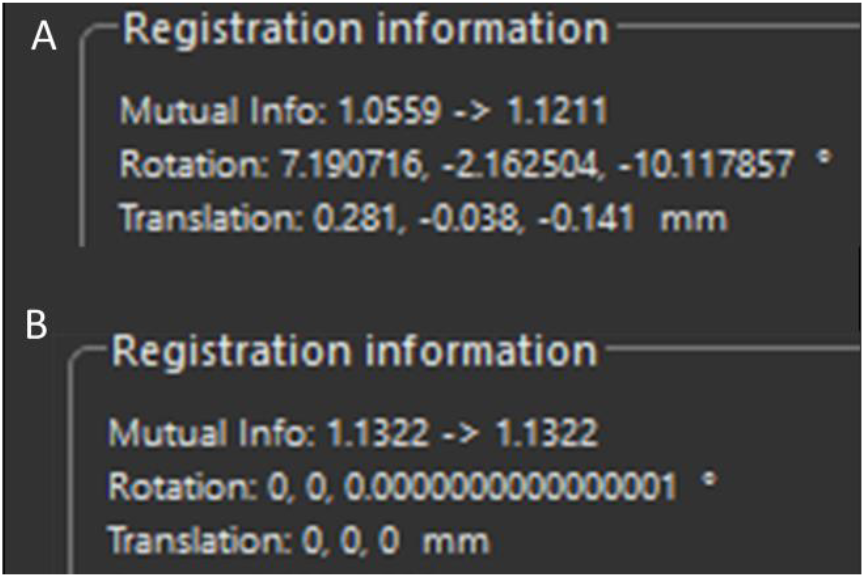
Registration information. A) Registration information following the first coarse registration step. B) Targeted registration information after the coarse, intermediate, and fine registration steps.

**Figure 6:**
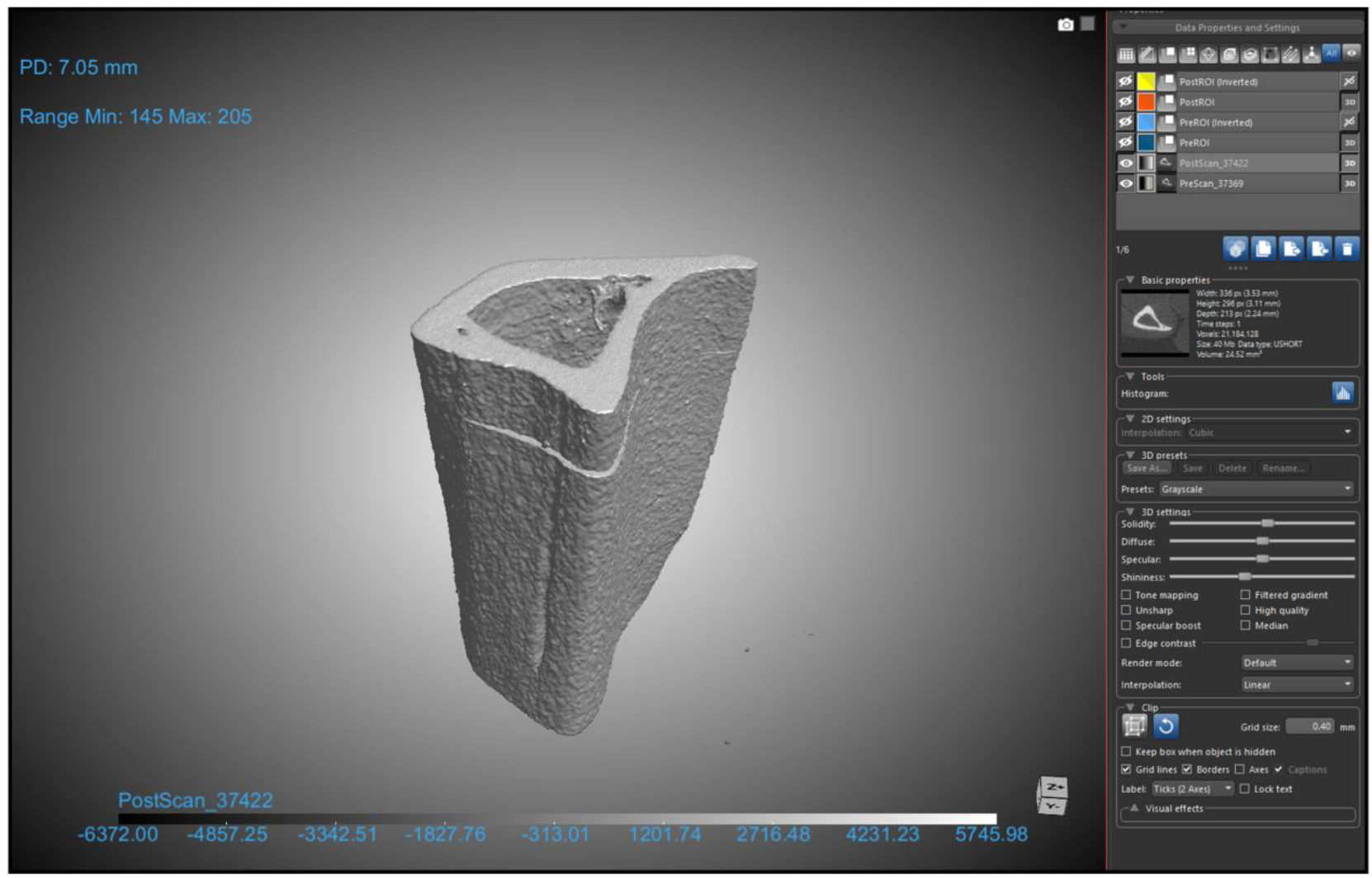
Dragonfly view of the registered PreScan and PostScan.

### Optional: Automated image registration using a macro

Step 15 described the image registration approach but requires manually applying registration parameters multiple times to achieve optimal alignment. This optional step provides an automated macro that executes the full registration sequence. Manual verification of registration quality is still required.

16. Automated image registration using a macro
  a. Download the macro file named “Meslier_RegistrationMacro” from github^8^ (https://github.com/QMuentin/3D-digital-dynamic-histomorphometry)
  b. Place the macro file in the Dragonfly macro directory: C:\Users\*username*\AppData\Local\ORS\Dragonfly2024.1\pythonUserExtensions\Macros *Note: Replace “username” with your own username. The directory path may vary depending on the location of the Dragonfly folder on your workstation. If the AppData folder is difficult to find, use Win+R, type %LocalAppData% and hit the “enter” key*.
  c. Restart Dragonfly if needed (Make sure to save your ORS session)
  d. After completing Steps 1–14, select Utilities → Macro Player
  e. From the drop-down menu, select “Meslier_RegistrationMacro”
  f. Click the “Play Next Step” button in the left bottom corner of the Macro Player to start the macro
  g. The macro will pause and prompt you to select the fixed and moving datasets.
  h. Select from the drop-down menus: PreScan_ScanNbr as the fixed channel and PostScan_ScanNbr as the moving channel
  i. Click “Play All Steps” to resume the macro. The automated registration will now run to completion
  j. After the macro finishes, manually repeat Step 15.h to verify that registration was successful. If the mutual information, rotation, or translation parameters are not satisfactory, perform manual refinement as described in Troubleshooting 4

*Note: See Troubleshooting 4*

### Create registered masks

#### Timing: < 5 min

17. Create Masks of registered scans (Fig.7)
  a. Select PreScan_ScanNbr to highlight it
  b. In the segment tab check the box “Define Range”
    1. Threshold should be the same as the Upper Otsu used at Step 12.c. We do not recommend clicking upper Otsu several times or playing with the value unless it is obviously off
  c. Click “Add to New” to create a new mask of the registered PreScan
  d. Rename the new mask as “PreBone”
  e. Smooth using the following parameters:
    1. Shape : Select “Circle”
    2. Dimensionality: Select “2D (Z)”
    3. Kernel size: Select “3”
  f. Repeat step 15 for PostScan_ScanNbr
    1. Adjust name of new mask to “PostBone”

**Figure 7:**
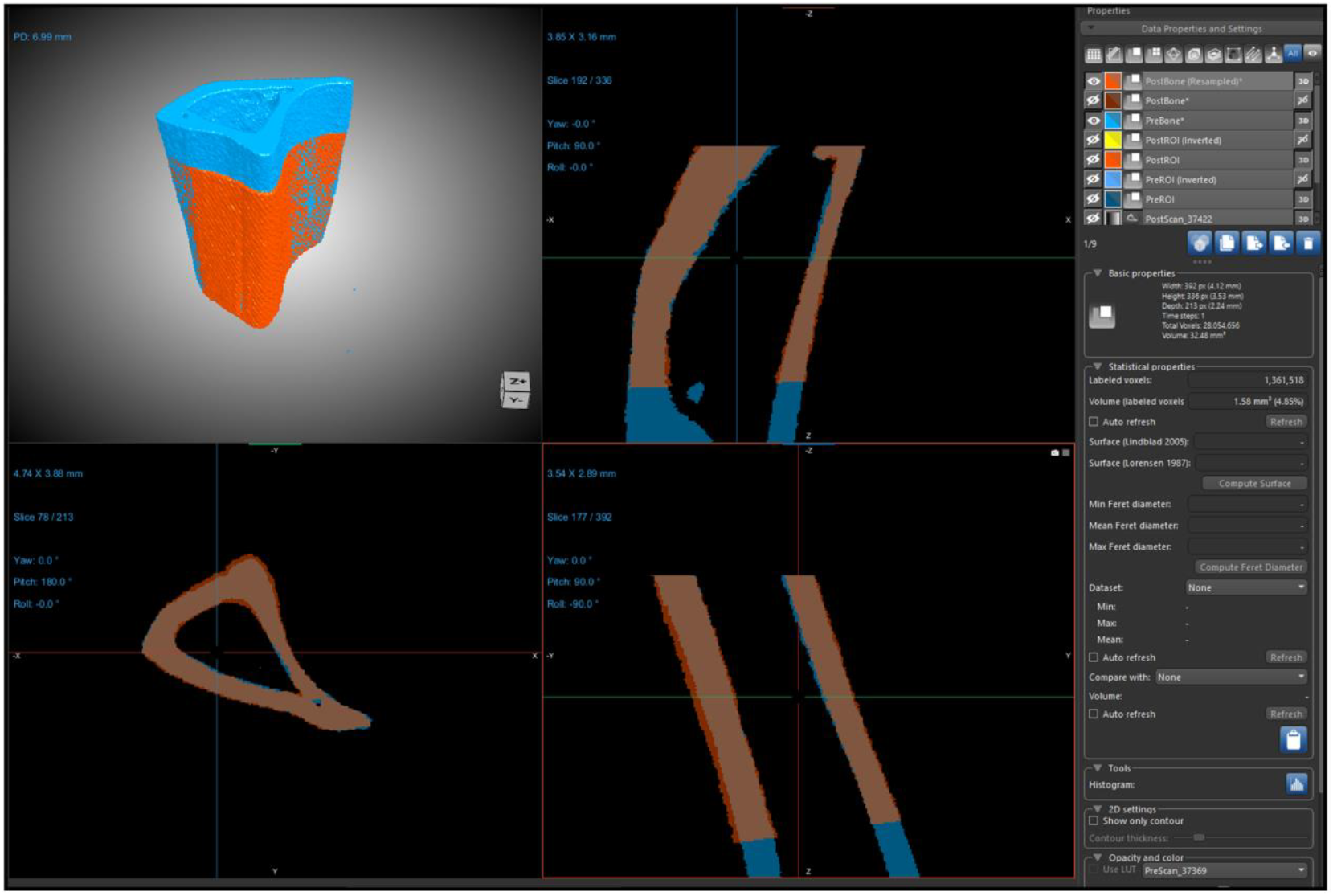
Dragonfly view of the registered Masks (PreBone and PostBone)

### Resample and Crop registered masks

#### Timing: < 5 min

Because the PostBone mask was generated in a different image space, it must be resampled to the PreScan_ScanNbr geometry to allow accurate comparison and further analysis.

18. Resample geometry of the PostBone maks.
  a. Right click on mask “PostBone”
  b. Go to “Modify and Transform”
  c. Click “Resample Geometry”
  d. Select “PreScan_ScanNbr” data set in the drop-down menu
  e. Click “OK”
    1. It will create a new mask named: “PostBone (Resampled)”
19. Crop overlapping masks and remove any non-overalping region of the registration. *Note: non-overlapping registration may happen because the region scanned during the Pre- and Post-scans were slightly different*.
  a. Make visible the following: “PreBone”, “PostBone (Resampled)”, and PreScan_ScanNbr
  b. Select PreScan_ScanNbr to highlight it. This is to guarantee that the slice number corresponds to the PreScan_ScanNbr
  c. Scroll through the transverse view to find the beginning and end slice where both “PreBone” and “PostBone (Resampled)” overlap
  d. Right click on “PostBone (Resampled)”
  e. Go to “Modify and Transform”
  f. Select “Crop”
  g. Enter the start and end z-slice identified in Step 19.c
  h. Check the box “Create New”
  i. Check the box “Apply to other”
  j. Click “Apply”
  k. Select only “PreBone” and “PostBone (Resampled)”
  l. Click “Ok”
  m. Click “Close”

### Remove artifacts

#### Timing: < 5min

Some scan artifacts may have intensity values similar to bone and can therefore be included during mask creation or thresholding.

20. Delete small pixels / artifacts. This step retains only the largest connected pixel region (representing the bone) and remove any small, isolated pixel clusters or artifacts.
  a. Right click on “PreBone (Cropped)”
  b. Select “Refine Region of Interest”
  c. Select “Process Islands”
  d. Select “Keep by largest 26 connected”
  e. Enter “1” in the pop-up menu
  f. Click “OK”

### Definition of bone formation, resorption, and quiescence

#### Timing: < 5min

Following image registration, Boolean operations are applied to the aligned datasets to define regions of bone formation, resorption, and quiescence.

21. Calculate formation, resorption, quiescent (Fig.8)
  a. Click “PreBone (Cropped)”
  b. Ctrl+click on “PostBone (Resampled) (Cropped)”
  c. Select correct task under Boolean Operations
    1. “Intersect”: Quiescent Bone
    2. “A-B”: Resorption
  d. Click on “PostBone (Resampled) (Cropped)” and ctrl+click on “PreBone (Cropped)”
  e. Click on “A-B”: Formation

**Figure 8:**
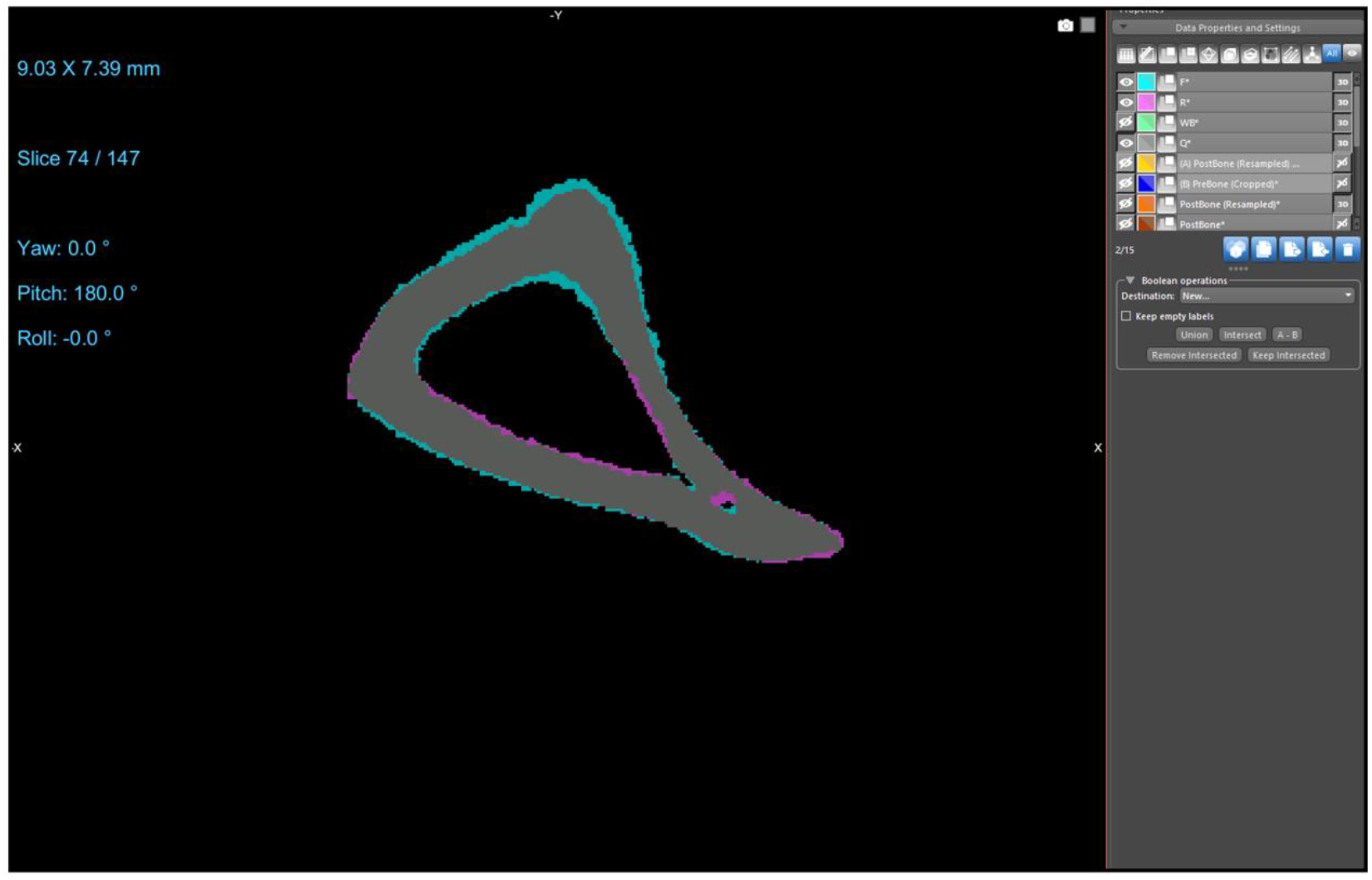
Dragonfly view showing the formation, resorption, and quiescent regions in mechanically loaded mouse tibia midshaft. Formation (cyan), resorption (magenta), and quiescent (gray) regions in a mouse tibia midshaft.

### Optional: Remove tibial ridge

#### Timing: 5 min

*Note: Depending on the research question, the tibial ridge region may not be relevant for quantification. Removing this region from the formation and resorption datasets at this stage reduces the number of processing steps required before surface-specific identification*.

22. Remove tibial ridge from the formation, resorption, and quiescent masks
  a. Select the quiescent mask “Q”
  b. In the “ROI painter”, make you way to the “2d view tools subsection
  c. Check the box “multi-slice”
  d. Select the squared paint brush
  e. In the drop-down menu, select “Full”
  f. In the transverse view, erase the part of the mask corresponding to the tibial ridge
    1. Start at the first slice of the mask
    2. Use “Ctrl+Scroll” to change the size of the brush
    3. Position the brush on the tip of the tibial ridge and use “shift+click” to remove it from the mask.
    4. Scroll through the stack until the mask reappears
    5. Repeat step 22.f.3 until you reach the top of the stack
  g. Repeat step 22 for masks “F”, and “R”

### Definition of surface specific formation and resorption

#### Timing: 5 min

This step allows for surface specific quantification as some experimental interventions may have effects on the periosteal but not the endosteal surface (Fig.9).

23. Define marrow at Pre timepoint
  a. Select the mask: “PreBone”
  b. Right click and select “Modify and Tranform”
  c. Select “Invert…”
  d. Check the box “Invert values” and the box “Create new”
  e. Right click on the newly created “PreBone (Inverted)”
  f. Select “Refine Region of Interest” and “Process Islands”
  g. Select “Remove by largest (26-connected)”
  h. Rename: “Marrow Pre” *Note: See Troubleshooting 5*
24. Define Marrow at Post timepoint
  a. Repeat step 21 for the mask “PostBone”
  b. Rename: “Marrow Post”
25. Define Endocortical surface (Fig.9)
  a. Select “Marrow Pre” and Ctrl+Click on “Marrow Post”
  b. Use Boolean operation “A-B”
  c. Rename “Endo_F” for endosteal formation
  d. Click “Ok”
  e. Select “Marrow Post” and Ctrl+Click on “Marrow Pre”
  f. Use Boolean operation “A-B”
  g. Rename “Endo_R” for endosteal resorption
  h. Click “Ok”
26. Define Periosteal parameters (Fig.9)
  a. Select mask “F” and Ctrl+Click on “Endo_F”
  b. Use Boolean operation “A-B”
  c. Rename to “Perio_F” for periosteal formation
  d. Click “Ok”
  e. Select mask “R” and Ctrl+Click on “Endo_R”
  f. Use Boolean operation “A-B”
  g. Rename to “Perio_R” for periosteal resorption
  h. Click “Ok”
27. Record volumes of each region in spreadsheet document
  a. Click on the mask of interest
  b. Under “Statistical Properties” highlight “Volume” and copy to spreadsheet
  c. Record: F, R, Endo_R, Endo_F, Perio_F, Perio_R, Q, and PreBone
28. Save ORS Session

**Figure 9:**
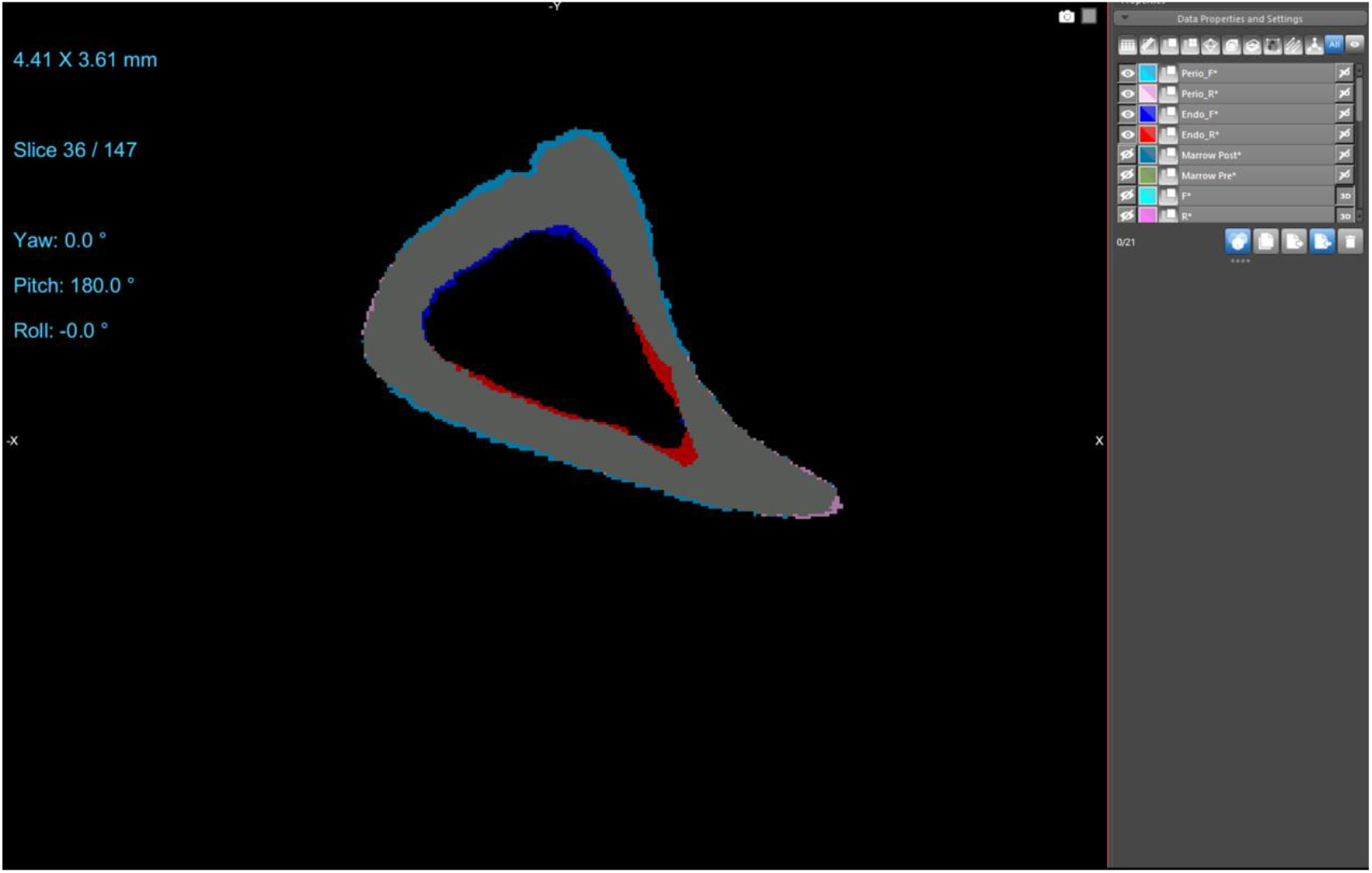
Definition of surface specific formation and resorption in a mechanically loaded mouse tibia midshaft. Periosteal formation: light blue. Periosteal resorption: pink. Endosteal formation: dark blue. Endosteal resorption: red.

This concludes with the main registration protocol. This part of the step-by-step protocol allows users to accurately register Pre- and Post-scans and gives access to quantification formation, resorption and quiescence from periosteal and endosteal surface. For results report, we recommend calculating the ratios of formed bone (Or resorbed bone) over PreBone as shown in the expected outcomes (Fig.14)

### Optional: MATLAB rendering and extra parameters quantification

The protocol described above enables straightforward, GUI-guided registration and quantification of bone scans. The quantification approach has been validated (see *Expected Outcomes*). In the following steps, we provide optional 3D rendering and additional parameter quantification. These optional outputs include 3D mineralizing and eroding surfaces, as well as 3D bone formation and resorption rates, providing complementary information on both the proportion of bone surface actively remodeling and the amount of bone formed/resorbed over the experimental period. Such measurements can help further characterize spatial remodeling patterns and improve interpretation of longitudinal bone adaptation.

These analyses rely on simple scripts and require only basic coding experience.

29. Export binary files from Dragonfly
  a. In Dragonfly, right click on the mask to export (e.g: “F”)
  b. Select “Export”
  c. Select “ROI as binary”
  d. Choose your directory to export the binary file
  e. Select “tiff” as file format
  f. Check the box “Output as one file”
  g. Repeat for masks: R, Q, WB, PreBone, Endo_F, Endo_R, Perio_F, and Perio_R *Note: if you performed option step 22 to remove the tip of the tibial ridge you can export those cropped masked as well*.
30. Prepare your experimental folder (Fig.10A)
  a. Create a subfolder “Samples”
  b. In “Samples”, create a folder and rename it with your sample number (e.g “624”)
  c. In “551” create a folder named “Mid” for Midshaft (Fig.10B).
    1. Repeat for “Proxi” if you have Proximal datasets
  d. In “Mid”, copy-paste the Dragonfly binary masks generated in step 29.g
31. Download the script entitle “Meslier_3DDynamicHisto_QM.m” from GitHub Repository ^8^and open the file in MATLAB
32. Change directories in the first section of the script:
  a. Copy the directory where the samples folders are located (e.g: ‘C:/Users/username/Samples/’ ) *Note: Mac requires “/Users/” instead of “C:/Users/”)*
  b. Paste the new directory in line 7 (Fig.11)
  c. Adjust the Folder number Line 5 (e.g: “624”) (Fig.11). This is the name of the folder containing the exported binary mask files (.tiff).
  d. Adjust the number of days that separate Pre- and Post-scans, Line 10 (Fig.11)
  e. Adjust the region being analyzed (Midshaft, Distal, Proximal), Line 12 (Fig.11)
  f. Adjust the voxels size in XY and in Z, Line 19 and 20 (Fig.10)
33. Download the Excel file template named “Meslier_Results_3DDynamicHisto_template.xlsx” from the supplementals
34. Copy Excel file to your experiment folder (Fig.10)
35. Prepare the excel file for results export
  a. In Excel the file, first sheet, enter your group names and sample numbers. This will allow for automatic assignment of the results in the different sheets.
  b. Save the modified Excel file as a copy and name it “Meslier_Results_3DDynamicHisto_Copy.xlsx”
  c. Close excel file
36. In the MALTAB script, change directory of the excel file for results export, Line 21 (Fig.11)
37. Click Run to run the MATLAB code
38. Quality check: The script will automatically generate various figures for quality assessment (Fig.12)
39. Repeat step 35 for all your folder/sample names

**Figure 10:**
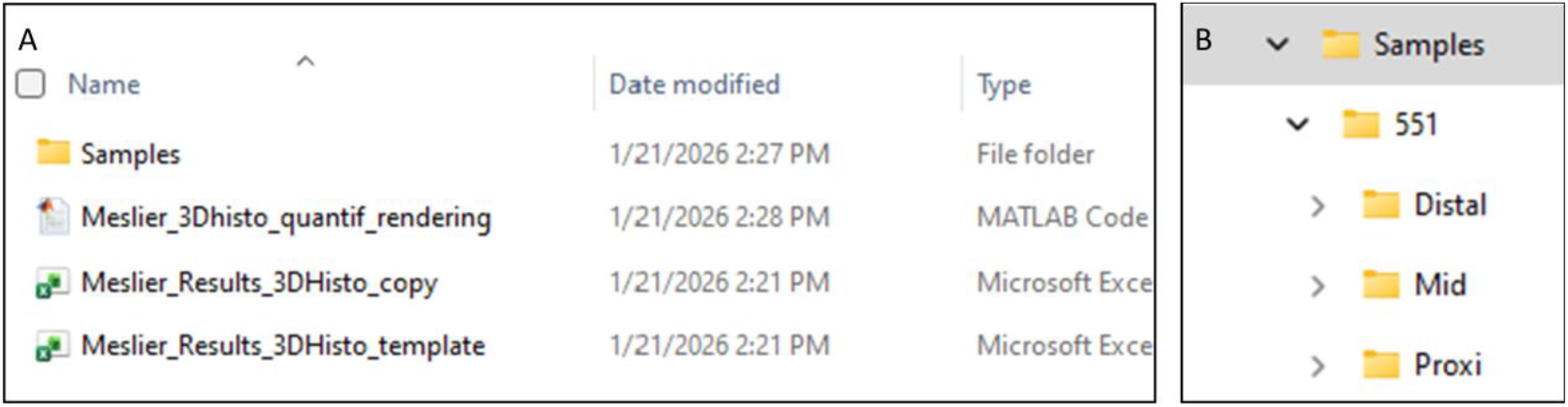
Optional MATLAB script set up A) Experimental folder organization. B) Samples subfolder organization

**Figure 11:**
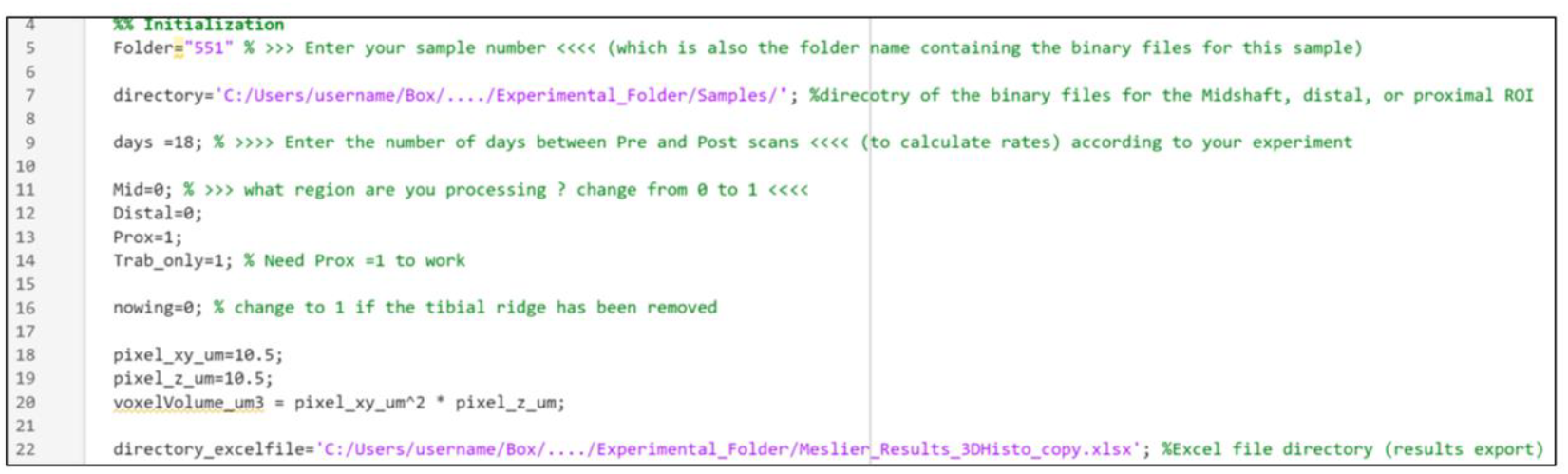
MATLAB script initialization section. The figure shows the lines of the code to be adjusted based on the user’ s directories, sample numbers, and region analyzed

**Figure 12:**
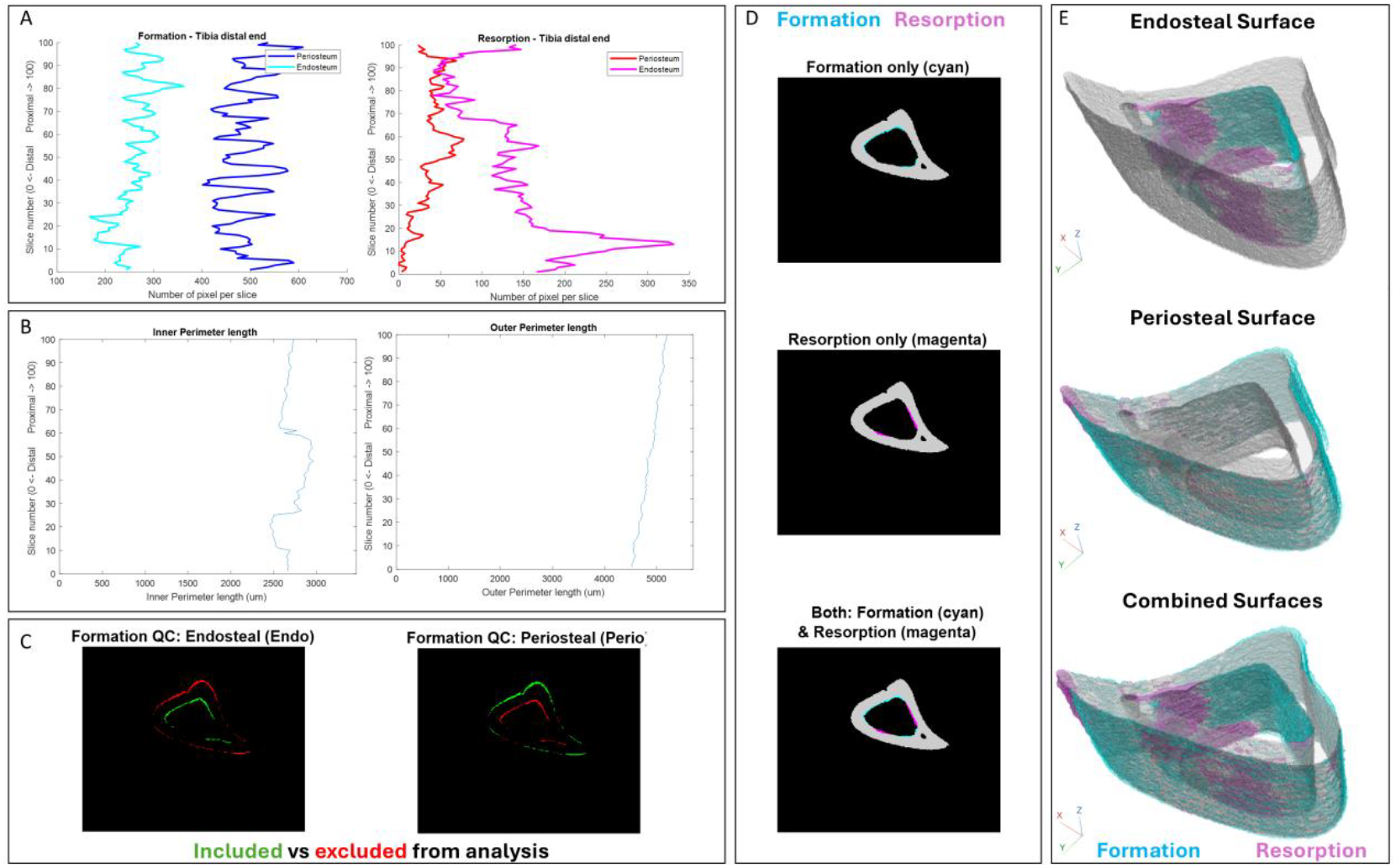
MATLAB-generated figures to facilitate evaluation of proper script execution and output quality. A) Number of pixels in the formation and resorption masks along the image stack. A larger number of pixels indicates a greater amount of detected bone formation or resorption in each slice. B) Perimeter length measured along the image stack. These plots allow verification that periosteal and endosteal surfaces are correctly identified. For tibiae, a smooth variation in perimeter length is expected across slices. Small irregularities may occur when trabecular bone partially closes the marrow cavity; however, sharp peaks restricted to a few slices indicate potential segmentation errors. C) Formation masks highlighting the endosteal and periosteal surfaces sequentially. Green regions indicate the surface currently included in the analysis, allowing verification that only the intended surface is quantified. D) Overlay of the cortical bone mask (“PreBone”, gray), formation (cyan), and resorption (magenta) on the endosteal surface. (E) Three-dimensional rendering of bone formation and resorption on the endosteal, periosteal, and combined surfaces.

*Note: For each sample, the code will also show the results in the MATLAB command window. Those results can be manually reported in a separate spreadsheet at the user convenience*.

### Optional: BoneJ quantification for Mineral Apposition and Resorption Rate parameters

These steps use masks generated by Dragonfly registration and the BoneJ plugin in Fiji to quantify the thickness of newly formed and resorbed bone, enabling calculation of 3D mineral apposition and resorption rates. These parameters add biological insight beyond surface measurements by determining the rate at which bone is deposited or removed at each remodeling site. This workflow does not require coding.

*Note: The user needs to download the BoneJ plugin and add it to fiji. Guidelines can be found on the BoneJ website*.

40. Export binary files from Dragonfly
  a. In Dragonfly, right click on the mask to export (e.g: “F” )
  b. Select “Export”
  c. Select “ROI as binary”
  d. Choose your directory to export the binary file
  e. Select “tiff” as file format
  f. Check the box “Output as one file”
  g. Repeat for masks: R, Q, Endo_F, Endo_R, Perio_F, and Perio_R
41. In Fiji, import one of the binary masks (e.g. the formation mask “F”)
42. Select “Plugins”
43. Select “BoneJ” and “Thickness”
44. Check the box “Trabecular thickness”, “Show thickness maps”, “Mask thickness maps”
45. Click “OK” *Note: Although it says, “Trabecular thickness”, the method will quantify the thickness of the regions of bone formation and resorption around the cortical bone*.
46. Assess the quality of the quantification by inspecting the thickness map, which displays thick bone regions in light yellow and thin regions in purple (Fig. 13)
47. A results table will automatically appear and show the mean thickness. Report results in a separate Excel file^3^
48. Divide the mean thickness results by the number of days between the pre-intervention scan and post-intervention scans to quantify the 3D mineral-apposition rate^3^
49. Repeat steps 39 to 47 for resorption masks to quantify the 3D mineral-resorption rate

**Figure 13:**
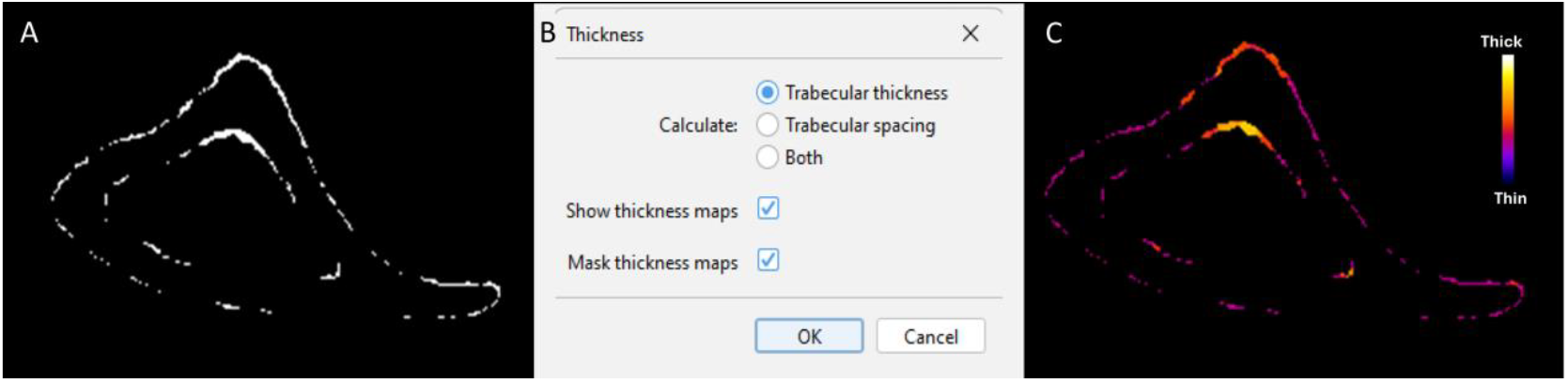
BoneJ quantification. A) Input binary mask (“F”) showing the region of bone formation in a mouse tibia midshaft. B) BoneJ “Thickness” menu. C) Resulting thickness map from BoneJ analysis.

## Results

Using this protocol, users can expect to obtain accurately registered pre- and post-scans of bone µCT datasets. Successful registration enables the identification and quantification of bone formation, resorption, and quiescent regions on both the endosteal and periosteal surfaces of trabecular and cortical bones. The protocol produces surface-specific and volume-based outputs, including measurements of newly formed and resorbed bone volumes that can be compared across experimental group (Table 4)

**Table 4:**
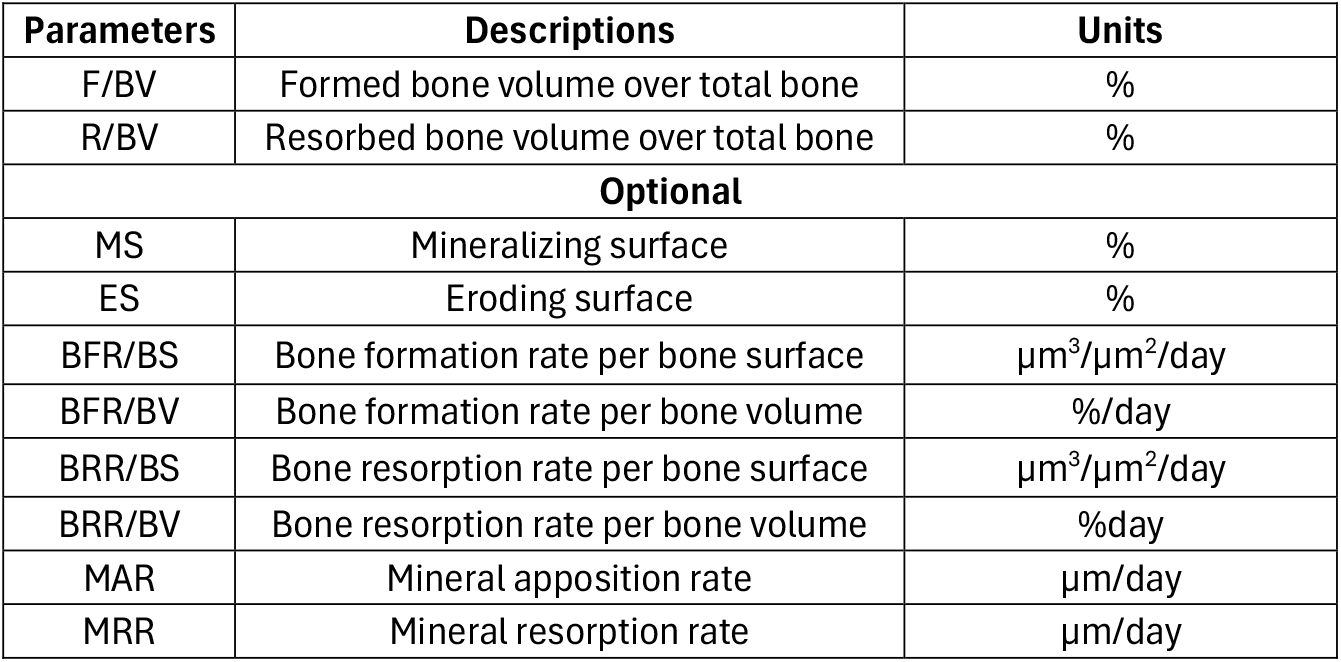
Expected parameters quantification using this protocol.

| Parameters | Descriptions | Units |
| --- | --- | --- |
| F/BV | Formed bone volume over total bone | % |
| R/BV | Resorbed bone volume over total bone | % |
| **Optional** |  |  |
| MS | Mineralizing surface | % |
| ES | Eroding surface | % |
| BFR/BS | Bone formation rate per bone surface | $\mu\text{m}^3/\mu\text{m}^2/\text{day}$ |
| BFR/BV | Bone formation rate per bone volume | %/day |
| BRR/BS | Bone resorption rate per bone surface | $\mu\text{m}^3/\mu\text{m}^2/\text{day}$ |
| BRR/BV | Bone resorption rate per bone volume | %day |
| MAR | Mineral apposition rate | $\mu\text{m}/\text{day}$ |
| MRR | Mineral resorption rate | $\mu\text{m}/\text{day}$ |

To validate the data generated by this protocol, periosteal bone formation quantified from registered µCT scans was compared with periosteal bone formation rates measured by 2D fluorochrome-based dynamic histomorphometry (Fig.14), a widely accepted gold-standard method. This comparison demonstrates that the protocol yields biologically meaningful and quantitatively reliable measures of bone formation suitable for longitudinal and group-based analyses. Proximal and distal tibia registrations by µCT were qualitatively compared with regions of calcein and alizarin labeling by dynamic histomorphometry (Fig. 15) and showed concordant regions of bone formation (yellow arrows). Areas lacking fluorescent labels in histomorphometry were consistent with regions of bone resorption identified through µCT registration (white triangles).

**Figure 14:**
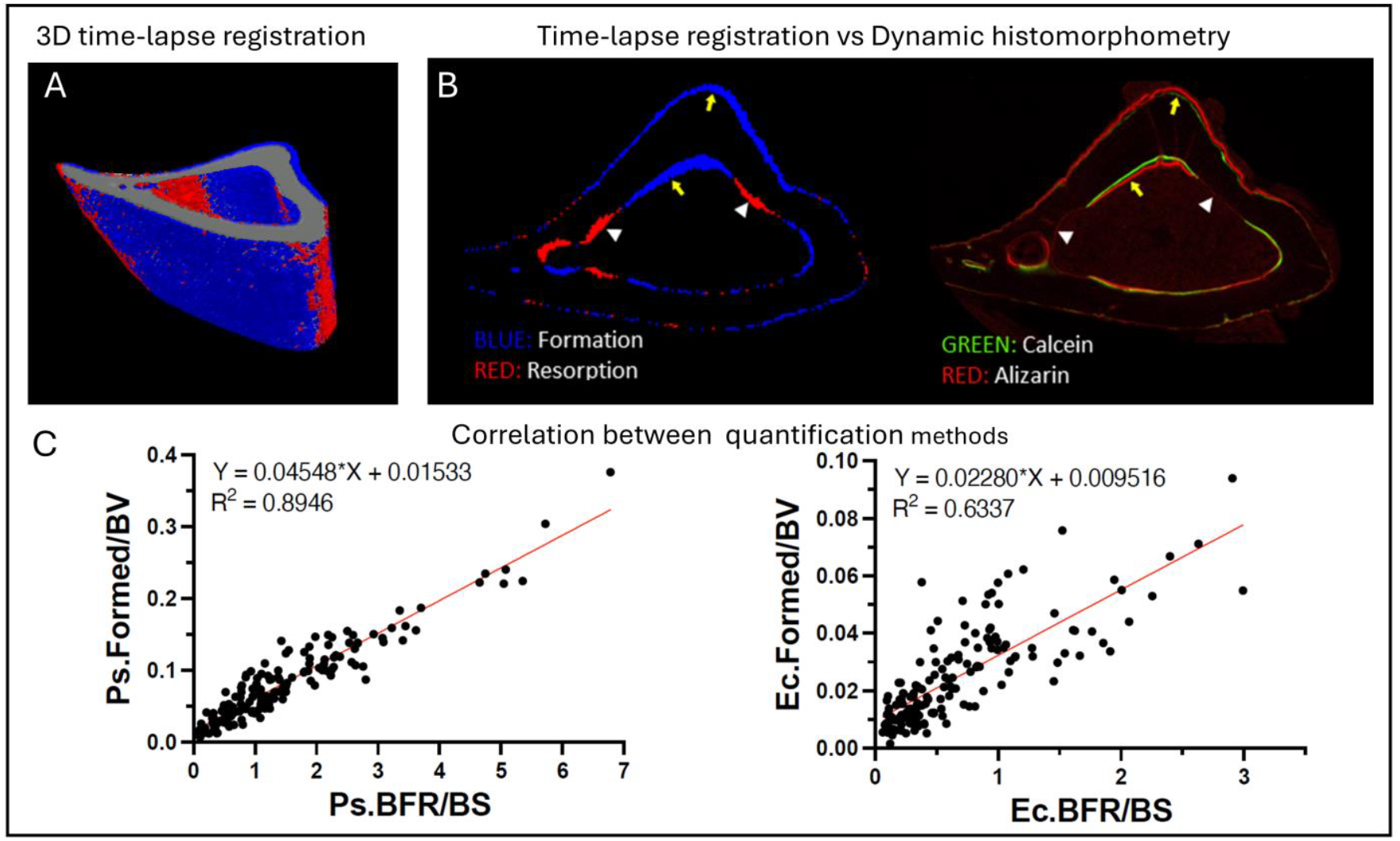
Validation of time-lapse registration in mouse tibial midshaft µCT scans. A) Three-dimensional time-lapse registration of mouse tibial midshaft µCT scans in Dragonfly (Blue = Formation. Red = Resorption). B) Comparison between time-lapse µCT-based registration and dynamic histomorphometry. Regions of bone formation are shown in blue in the time-lapse registration and are visualized by calcein green and alizarin red labeling in dynamic histomorphometry. Yellow arrows indicate corresponding regions of bone formation identified by both methods, while white triangles indicate regions of bone resorption. C) Correlation between methods. Periosteal (Ps.Formed/BV) and endocortical (Ec.Formed/BV) bone formation normalized to bone volume, quantified from Dragonfly longitudinal µCT registration (y-axis), correlates with relative periosteal (Ps.BFR/BS) and endocortical (Ec.BFR/BS) bone formation rate measured by dynamic histomorphometry (x-axis).

**Figure 15:**
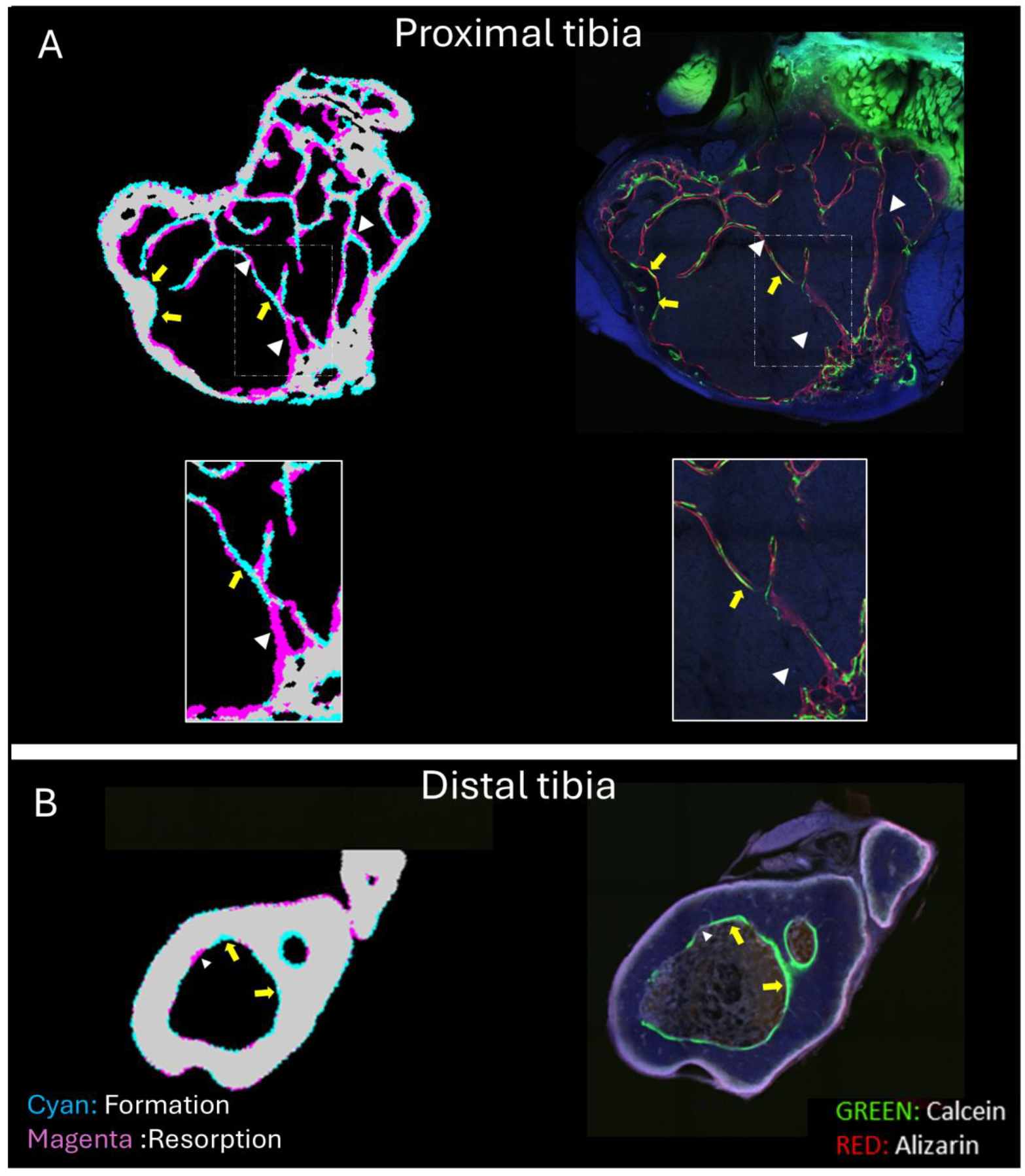
Comparison between time-lapses *in vivo* microCT registration and dynamic histomorphometry for proximal and distal tibia. A) Proximal tibia registration and dynamic histomorphometry comparison. B) Distal tibia registration and histomorphometry comparison.

A key advantage of this technique over traditional histologic methods is that it enables visualization and quantification of three-dimensional bone formation and resorption across a substantially larger region of interest than is possible with a single histologic section. This approach allows surface-specific quantification at the endosteal, periosteal, or combined surfaces (Fig.9,14C). In addition, an optional MATLAB-based rendering workflow enables enhanced three-dimensional visualization of formation and resorption volumes (Fig. 16).

**Figure 16:**
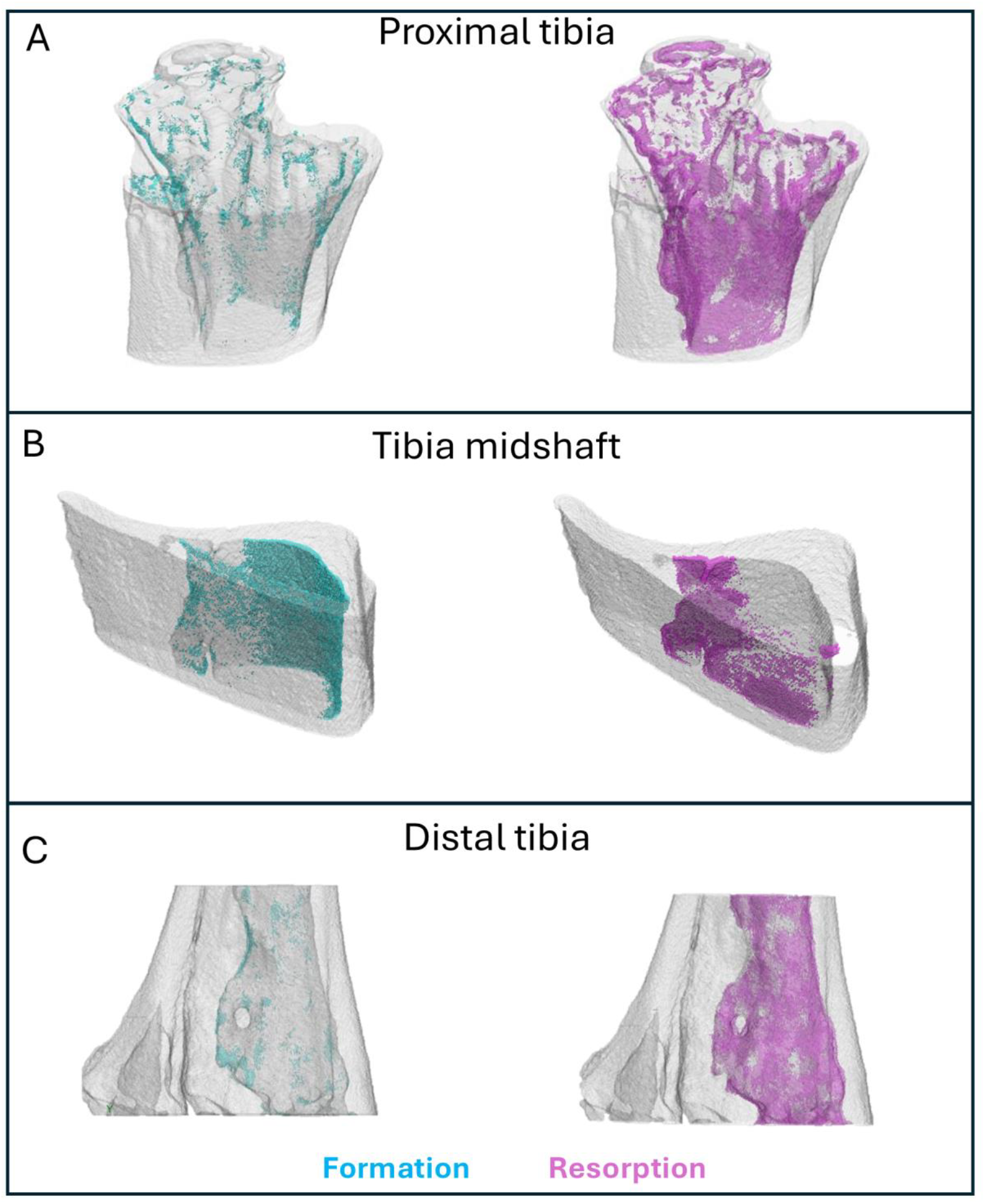
MATLAB 3D rendering of bone formation and resorption on the endosteal surface. A) In the proximal tibia. B) Tibia Midshaft. C) Distal tibia.

**Figure 17:**
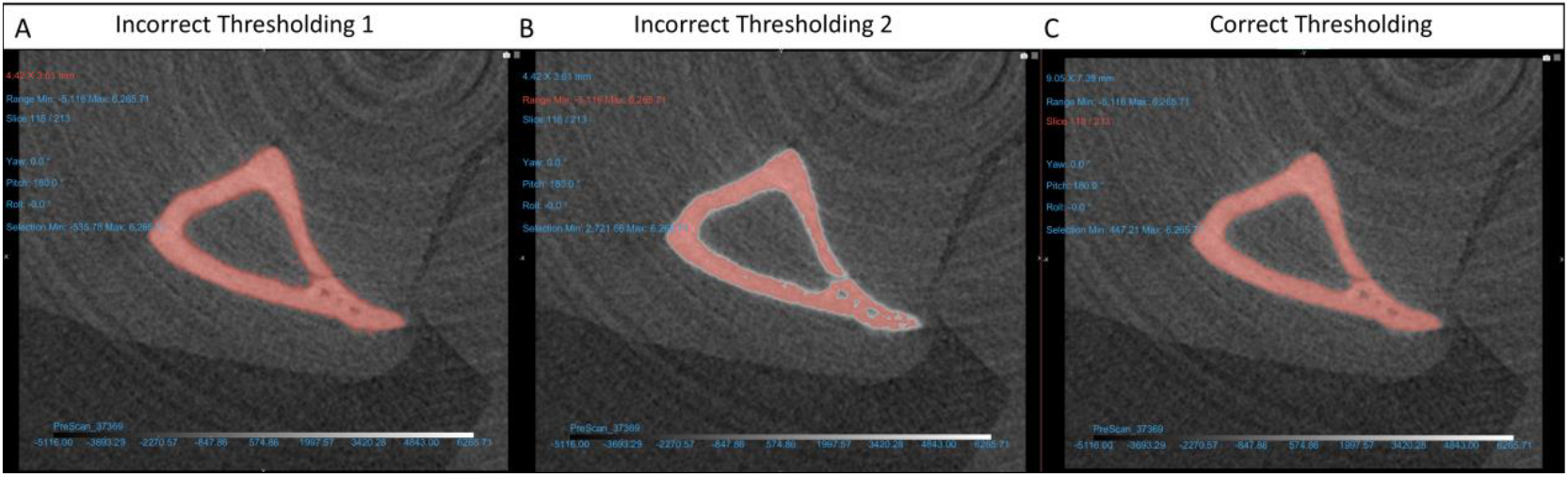
Bone thresholding in Dragonfly. A) Thresholding includes bone and some of the background. B) Thresholding does not include all the cortical bone. C) Correct thresholding using upper Otsu function.

In a recent study, we applied this protocol to investigate the effects of uniaxial tibial loading on bone formation and resorption in mice with a conditional deletion of the citrate transporter *Slc13a5*, compared to control littermates. ^9^.

### Limitations

This protocol relies on accurate registration of longitudinal µCT scans; registration quality may be reduced in datasets with substantial differences in positioning, image noise, motion artifacts, or anatomical alterations between time points.

Although the automated registration macro facilitates alignment, challenging scans may require manual refinement, and incomplete registration can affect the accuracy of downstream Boolean operations used to identify formation, resorption, and quiescent regions.

Threshold-based mask generation represents an additional limitation, as imaging artifacts or non-bone structures with intensity values similar to mineralized tissue may be inadvertently included. Such artifacts can influence surface-specific and volumetric measurements if not corrected through manual inspection and cleanup. Variability in scan quality, voxel size, and acquisition parameters across instruments or studies may also affect quantitative comparability. In particular, scan resolution defines the minimum magnitude of bone formation or resorption that can be interpreted as biologically meaningful; changes below this threshold cannot be interpreted with confidence.

Bones with symmetrical structure around all axis (Femur mid diaphysis for example) represent more challenging registration due to the lack of asymmetric anatomical features (landmarks). Including an anatomical landmark such as the trochanter in the ROI will facilitate the registration.

This protocol relies on repeated *in vivo* µCT scans which may have detrimental effects on bone structure depending on the level of the radiation, frequency of scanning, and recovery time between scanning session ^10–12^. Those potential adverse effects should be considered in experimental design.

Finally, this protocol quantifies mineralized tissue changes detected by µCT and does not directly capture cellular activity or early, non-mineralized bone formation (osteoid). As a result, measurements reflect net mineralized bone dynamics rather than underlying osteoblast or osteoclast activity. Validation against histological or dynamic histomorphometry approaches is therefore recommended when biological interpretation requires cellular-level resolution.

## Troubleshooting

### Problem 1: Upper Otsu and thresholding

If you click several times of the upper Otsu, this might result in incorrect thresholding (Fig.17B). In addition, sometimes some background artifacts might be included in the upper Ostu thresholding.

#### Potential solution

If this happens you can either delete and reload the Dicom file, or you can use the value that you noted in Step 12.d to manually adjust the threshold. If the upper Otsu includes a lot of background artifacts you can manually adjust the threshold to a correct thresholding (Fig.17C). Or you can use the upper Otsu thresholding and manual remove artifact from the mask using the ROI painter.

### Problem 2: Align Centroids

If the Pre- and Post-scans have a very different initial orientation, step 14 might fail.

#### Potential solution

If the “Align centroids” step fail, a first simple manual alignment can help. In the right panel, select the Post-Scan in the “Data Properties and Settings” tab. On the left panel, under “Main”-> “Translate/Rotate, select “Displace”. Click on the transverse view to make the displacement commands appear. Use the translation and rotation commands to roughly align the two scans. The rough alignment can be checked via the 3D view.

### Problem 3: Image registration plugin

After you selected Utilities -> Open Plugins -> Image registration and that the new image registration window popped up. Sometimes the software does not allow manual changes to image registration values. This is a glitch.

#### Potential solutio

If this happens, switch “fixed” and “Moving” images. Then switch them back again so that the PreScan_ScanNbr is the Fixed image and the PostScan_ScanNbr is the moving image. This should resolve the issue.

#### Problem 4: Automated image registration macro

After completing optional Step 16 (automated image registration using the macro), Step 15.h must be performed manually to verify registration quality. The automated macro may fail to achieve optimal alignment in cases where scans are more challenging to register. In such cases, discrepancies in the mutual information metric indicate incomplete registration and require manual correction.

#### Potential solution

Complete the automated registration by manually running the image registration plugin starting at Step 15.h, iterating as needed until the “mutual information”, “rotation” and “translation” parameters reach acceptable values.

### Problem 5: Definition of marrow space

If the bone marrow space is connected to the outer space of the bone (via a blood vessel for example) then the “Remove by Largest” (Step 22.g) will remove both the outer space and the marrow area.

#### Potential solution

Manually remove all connection between the bone marrow space and the outside space of the bone using the ROI painter erase (Fig.18). Use “single slice” if the connection is only in one image. Use Shift+Scroll to adjust the size of the brush. Use Shift+Click to remove the blood vessel from the mask.

**Figure 18:**
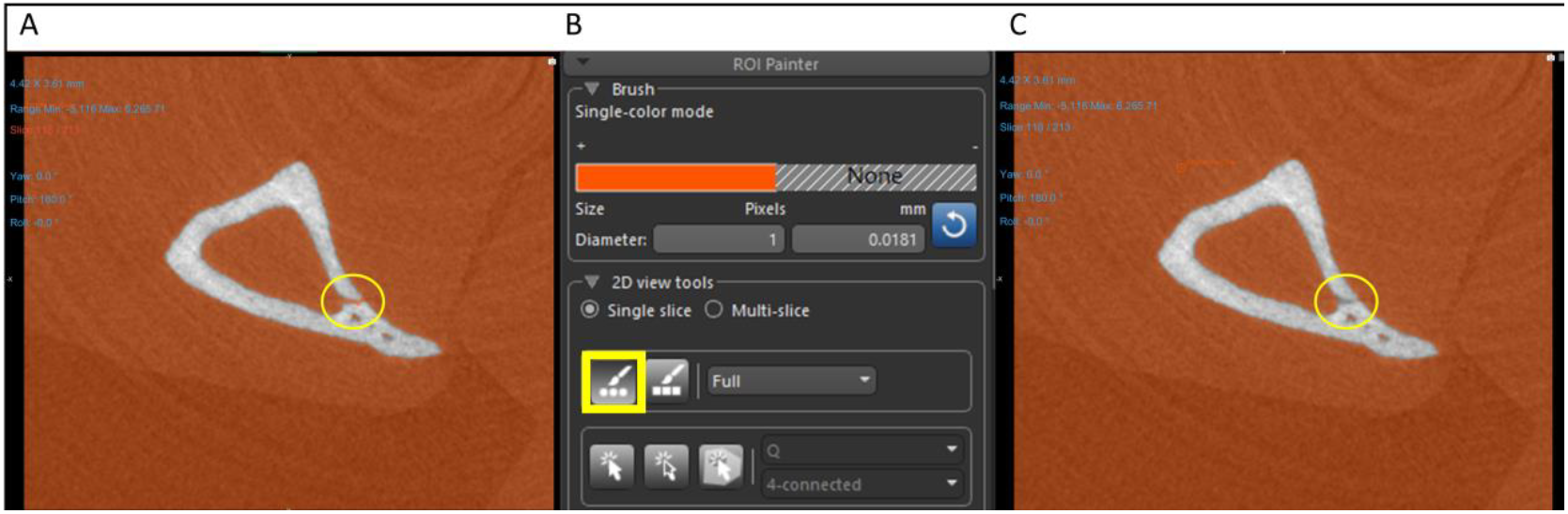
Marrow space definition. If the marrow space is connected via blood vessel (yellow circle), it needs to be manually disconnected using the ROI painter tools

## Resource availability

### Lead contact

Further information and requests for resources and reagents should be directed to and will be fulfilled by the lead contact, Dr. Silva

### Technical contact

Technical questions on executing this protocol should be directed to and will be answered by the technical contact, Dr. Meslier

### Materials availability

- This study did not generate new unique reagents.

### Data and code availability statement

- The Matlab code and Dragonfly macro generated during this study are available on GitHub: [3D-digital-dynamic-histomorphometry] (https://github.com/QMuentin/3D-digital-dynamic-histomorphometry)

## Author contributions

*Q.A.M: Writing original draft, Data curation, formal analysis, Methodology, Software, Visualization*

*N.M: Writing – Review & Editing, Data curation, formal analysis, Methodology, Software, Visualization*

*S.N.L: Writing – Review & Editing, Methodology, Software*

*E.L.S: Writing – Review & Editing, Funding acquisition, Supervision*

*M.J.S: Writing – Review & Editing, Funding acquisition, Supervision*

